# Molecular mechanism underlying the non-essentiality of Bub1 for the fidelity of chromosome segregation in human cells

**DOI:** 10.1101/2021.02.01.429225

**Authors:** Qinfu Chen, Miao Zhang, Xuan Pan, Linli Zhou, Haiyan Yan, Fangwei Wang

**Author notes:** Contributed equally (Qinfu Chen, Miao Zhang, Xuan Pan). To whom correspondence should be addressed: Fangwei Wang, Life Sciences Institute, Zhejiang University, Nano Building, Room 577, 866 Yuhangtang Road, Hangzhou, Zhejiang 310058, China.

## Abstract

The multi-task protein kinase Bub1 has long been considered important for chromosome alignment and spindle assembly checkpoint signaling during mitosis. However, recent studies provide surprising evidence that Bub1 may not be essential in human cells, with the underlying mechanism unknown. Here we show that Bub1 plays a redundant role with the non-essential CENP-U complex in recruiting Polo-like kinase 1 (Plk1) to the kinetochore. While disrupting either pathway of Plk1 recruitment does not affect the accuracy of whole chromosome segregation, loss of both pathways leads to a strong reduction in the kinetochore accumulation of Plk1 under a threshold level required for proper chromosome alignment and segregation. Thus, parallel recruitment of Plk1 to kinetochores by Bub1 and the CENP-U complex ensures high fidelity of mitotic chromosome segregation. This study may have implications for targeted treatment of cancer cells harboring mutations in either Bub1 or the CENP-U complex.

## INTRODUCTION

The precise segregation of chromosomes into daughter cells during mitotic division is essential for cellular and organismal viability. Error-free chromosome segregation during mitosis requires a functional spindle assembly checkpoint (SAC), which is a surveillance mechanism that delays anaphase onset until all the chromosomes are properly bi-oriented on the mitotic spindle (Musacchio, 2015). Metaphase chromosome bi-orientation depends on proper attachment of their kinetochores to spindle microtubules. Previous studies using siRNA-mediated knockdown demonstrate important roles for the evolutionarily conserved kinetochore protein Bub1 in SAC signaling and chromosome congression (Elowe, 2011, Johnson *et al*, 2004, London & Biggins, 2014b).

Bub1 localizes to the outer kinetochore during mitosis, which relies on Mps1-mediated phosphorylation of Knl1 at multiple conserved MELT motifs. Knl1 recruits Bub3-Bub1 complexes via the direct interaction of Bub3 with phospho-MELT motifs (London *et al*, 2012, Primorac *et al*, 2013, Shepperd *et al*, 2012, Vleugel *et al*, 2015, Yamagishi *et al*, 2012). Mps1 further phosphorylates Bub1 to promote its interaction with Mad1-Mad2 complexes, thereby producing a SAC signaling platform (London & Biggins, 2014a, Wenjin & Jianwei, 2017). The Bub1-Mad1 platform is thought to recruit Bub3, BubR1, Cdc20, and Mad2 to generate the SAC effector mitotic checkpoint complex (MCC). Bub1 also binds the Polo-box domain (PBD) of Plk1 and promotes Plk1 localization at kinetochores (Qi *et al*, 2006). The Bub1-Plk1 kinase complex promotes SAC signaling through phosphorylating Cdc20 (Jia *et al*, 2016, Tang *et al*, 2004a). Therefore, Bub1 plays a key role in the assembly of SAC proteins at the outer kinetochore (London & Biggins, 2014b). Besides, the BUB3-BUB1 complex is also reported to promote telomere DNA eplication (Li *et al*, 2018).

Moreover, kinetochore-localized Bub1 mediates histone H2A threonine 120 phosphorylation (H2ApT120) at mitotic centromeres, which recruits Shugoshin to protect centromeric cohesion (Kawashima *et al*, 2010, Kitajima *et al*, 2005, Kitajima *et al*, 2004, Liu *et al*, 2013a, Liu *et al*, 2015, Liu *et al*, 2013b, Tang *et al*, 2004b), and to recruit the Aurora B kinase-containing chromosomal passenger complex (CPC) that corrects erroneous kinetochore-microtubule (KT-MT) attachments (Tsukahara *et al*, 2010, Yamagishi *et al*, 2010). H2ApT120 also recruits DNA topoisomerase II α (TOP2A) to mitotic centromeres to promote the decatenation of centromeric DNA (Zhang *et al*, 2019c). Thus, Bub1 kinase activity plays a pivotal role in the assembly of the functional centromere (Boyarchuk *et al*, 2007).

Each of these demonstrated functions of Bub1 at the kinetochore-centromere region is considered to be important for mitosis. Indeed, Bub1 is essential in a number of model organisms including fission yeast and budding yeast. However, the essentiality of Bub1 in human cell lines was recently challenged by CRISPR/Cas9-mediated gene knockout in the diploid and non-transformed human RPE1 cells, as well as in human haploid HAP1 cells (Currie *et al*, 2018, Raaijmakers *et al*, 2018, Zhang *et al*, 2019a). These Bub1 knockout cells seemed to have almost fully functional SAC response. Subsequent studies reported that these Bub1 cell lines express residual Bub1 protein (Rodriguez-Rodriguez *et al*, 2018, Zhang *et al*, 2019b), which might be produced via nonsense-associated alternative splicing. A subsequent study succeeded in generating a full Bub1 knockout in HAP1 cells (Raaijmakers & Medema, 2019), indicating that HAP1 cells can survive in the complete loss of Bub1.

In addition, Zhang et al. used CRISPR/Cas9 and obtained multiple Bub1 knockout HeLa cell lines in which Bub1 was not detectable by immunoblotting (Zhang *et al*, 2019b). Based on mass spectrometry analysis, the authors estimated that 4% Bub1 was left in these Bub1 “knockout” cell lines. In these cells, chromosome alignment was delayed, but cells rarely entered anaphase with unaligned chromosomes due to a functional checkpoint. Moreover, these cell lines exhibited a normal checkpoint response to the spindle toxin nocodazole.

Collectively, these studies demonstrate that Bub1 can be fully knocked out in HAP1 cells (Raaijmakers & Medema, 2019), and that 96% loss of Bub1 protein does not affect the fidelity of chromosome segregation (Zhang *et al*, 2019b), raising the question of why Bub1 is largely dispensable for proper mitosis progression in human cells.

In this study, we knocked out Bub1 in HeLa cells using CRISPR/Cas9-mediated genome editing. We present data that the inner kinetochore CENP-U complex comprising CENP-O, CENP-P, CENP-Q and CENP-U, which is non-essential in most human cell lines (McKinley & Cheeseman, 2017, McKinley *et al*, 2015, Wang *et al*, 2015), is critically required to support accurate chromosome segregation in the absence of Bub1. Mechanistically, we show that CENP-U and Bub1 recruit Plk1 to the kinetochore redundantly to promote chromosome alignment, thereby ensuring the fidelity of mitotic chromosome segregation.

## RESULTS

### Knockout of Bub1 severely reduces kinetochore/centromere localization of Plk1, Bub3, Mad1, BubR1, TOP2A, Sgo1 and Aurora B

To analyze the role for Bub1 in mitosis, we attempted to knock out Bub1 in HeLa cells using CRISPR/Cas9. We obtained a number of clonal cell lines in which H2ApT120 was undetectable by immunofluorescence microscopy. Genome sequencing detected indels in clones 2D1, 2D4, 2B6 and 2C1 (Figure S1). Bub1 protein was analyzed by immunoblotting and immunofluorescence microscopy using two polyclonal antibodies (Taylor *et al*, 2001, Zhang *et al*, 2019c). Immunoblotting demonstrated various levels of Bub1 protein in these cell lines, among which clones 2B6 and 2C1 appeared to be Bub1 knockout cell lines (Figure 1A; see also in Figure 7C). Immunofluorescence microscopy did not detect kinetochore localization of Bub1 in these two clones (Figure 1B and 1C). Hereafter, clones 2B6 and 2C1 cells were referred to as ΔBub1 cells.

**Figure 1.**
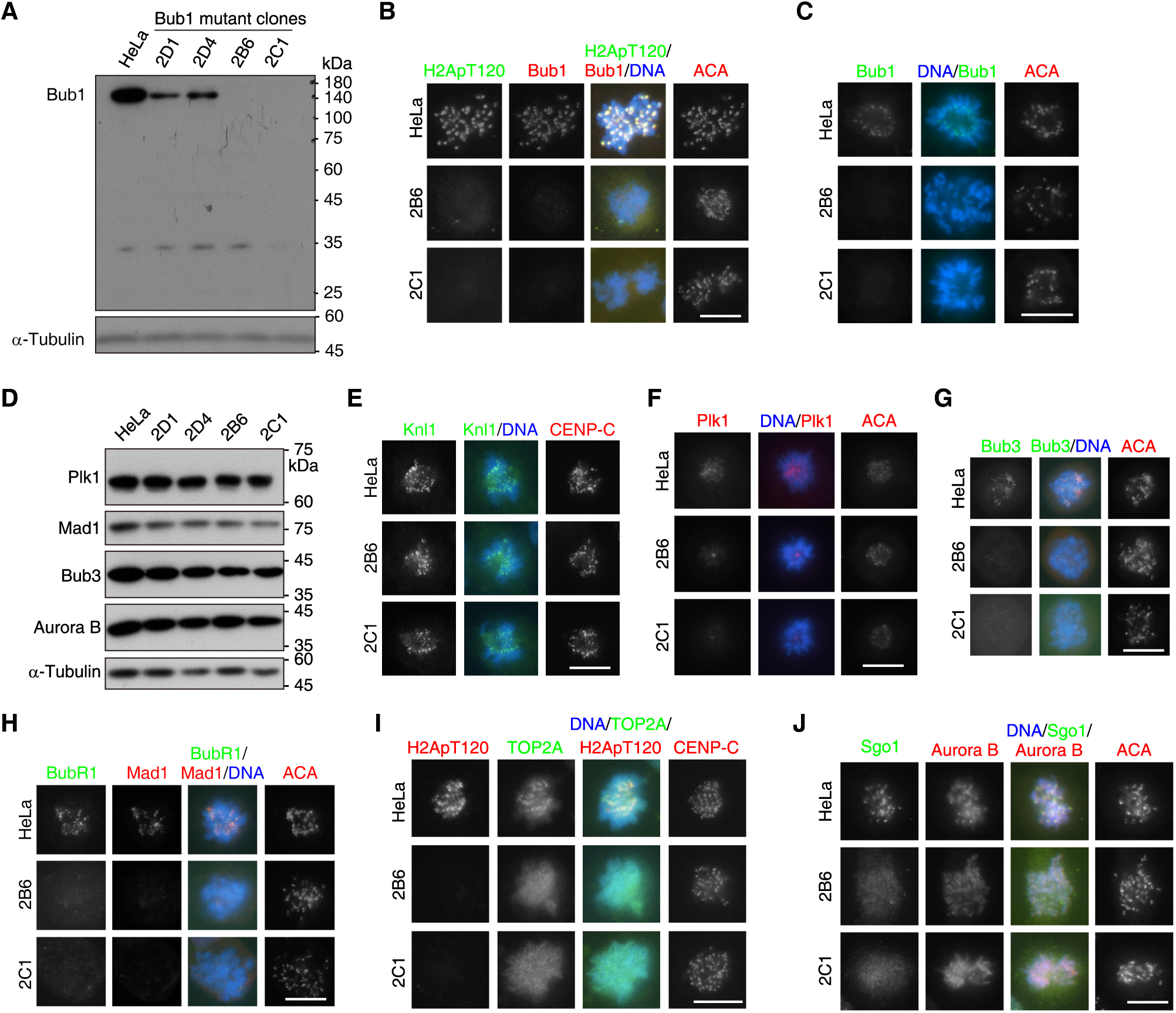
Knockout of Bub1 severely reduces kinetochore/centromere localization of Plk1, Bub3, Mad1, BubR1, TOP2A, Sgo1 and Aurora B. (A) HeLa cells and the indicated Bub1 mutant clones were treated with nocodazole for 14 h, then mitotic cells were collected and analyzed by immunoblotting with the α-Tubulin antibody and sheep anti-Bub1 polyclonal antibodies. (B) HeLa cells and the indicated ΔBub1 clones were immunostained with the anti-human centromere autoantibody (ACA), the H2ApT120 antibody and sheep polyclonal anti-Bub1 antibodies. DNA was stained by DAPI. (C) HeLa cells and the indicated ΔBub1 clones were immunostained with ACA and rabbit anti-Bub1 polyclonal antibodies. (D) HeLa and the indicated Bub1 mutant clones were treated with nocodazole for 14 h, then mitotic cells were collected. The samples were analyzed by immunoblotting with the indicated antibodies. (E) HeLa cells and the indicated ΔBub1 clones were immunostained with antibodies for Knl1 and CENP-C. (F) HeLa cells and the indicated ΔBub1 clones were treated with STLC for 5 h and then immunostained with ACA and the Plk1 antibody. (G and H) HeLa cells and the indicated ΔBub1 clones were treated with 20 ng/ml nocodazole for 4 h and then immunostained with ACA and antibodies either for Bub3 (G) or for BubR1 and Mad1 (H). (I and J) HeLa cells and the indicated ΔBub1 clones were immunostained either with antibodies for H2ApT120, TOP2A and CENP-C (I), or with ACA and antibodies for Sgo1 and Aurora B (J). Scale bars, 10 μm. See also Figure S1.

We next examined the effect of Bub1 loss on proteins whose localization at centromeres/kinetochores requires Bub1. Immunoblotting showed that the protein levels of Plk1, Bub3, Mad1 and Aurora B did not obviously change in ΔBub1 cells (Figure 1D). Immunofluorescence microscopy demonstrated that Knl1 localized normally to kinetochores in ΔBub1 cells (Figure 1E; see also in Figure 4D), indicating that the outer kinetochore was properly assembled. In contrast, kinetochore localization of Plk1 (Figure 1F), Bub3 (Figure 1G) and Mad1 (Figure 1H) was drastically reduced, which is in line with their direct interaction with Bub1 at the kinetochore. Moreover, consistent with the role for Bub3 in binding BubR1 (Johnson *et al*, 2004), the kinetochore localization of BubR1 was also reduced in ΔBub1 cells (Figure 1H).

In addition, due to the loss of H2ApT120, TOP2A and Sgo1 were no longer enriched at mitotic centromeres in ΔBub1 cells (Figure 1I and 1J). Aurora B was less concentrated at centromeres in ΔBub1 cells and, together with Sgo1, displayed diffused signals along the length of chromosomes (Figure 1J; see also in Figure 4C), which is in agreement with the role for Sgo1 in binding the CPC (Tsukahara *et al*, 2010).

Thus, the kinetochore/centromere localization of a number of important mitotic regulators is largely reduced in ΔBub1 cells, due to the loss of Bub1 or H2ApT120.

### Loss of Bub1 does not compromise chromosome segregation fidelity during unperturbed mitosis

We assessed the impact of Bub1 loss on chromosome segregation during otherwise unperturbed mitosis. Fluorescence microscopy of asynchronous cells revealed that the frequency of anaphases with lagging chromosomes was essentially unchanged in ΔBub1 cells compared to control HeLa cells (4.1% vs 2.5%) (see Figure 3C and 3F). Consistently, the percentage of pre-anaphase mitotic cells with misaligned chromosomes was only slightly higher in ΔBub1 cells (8.1%-9.6%) than in control HeLa cells (4.6%) (Figure 2A and 2B). Moreover, after treatment for 3 h with MG132, a proteasome inhibitor which induces metaphase arrest, around 95% of HeLa cells and ΔBub1 cells had completed metaphase chromosome alignment (Figure 2C).

**Figure 2.**
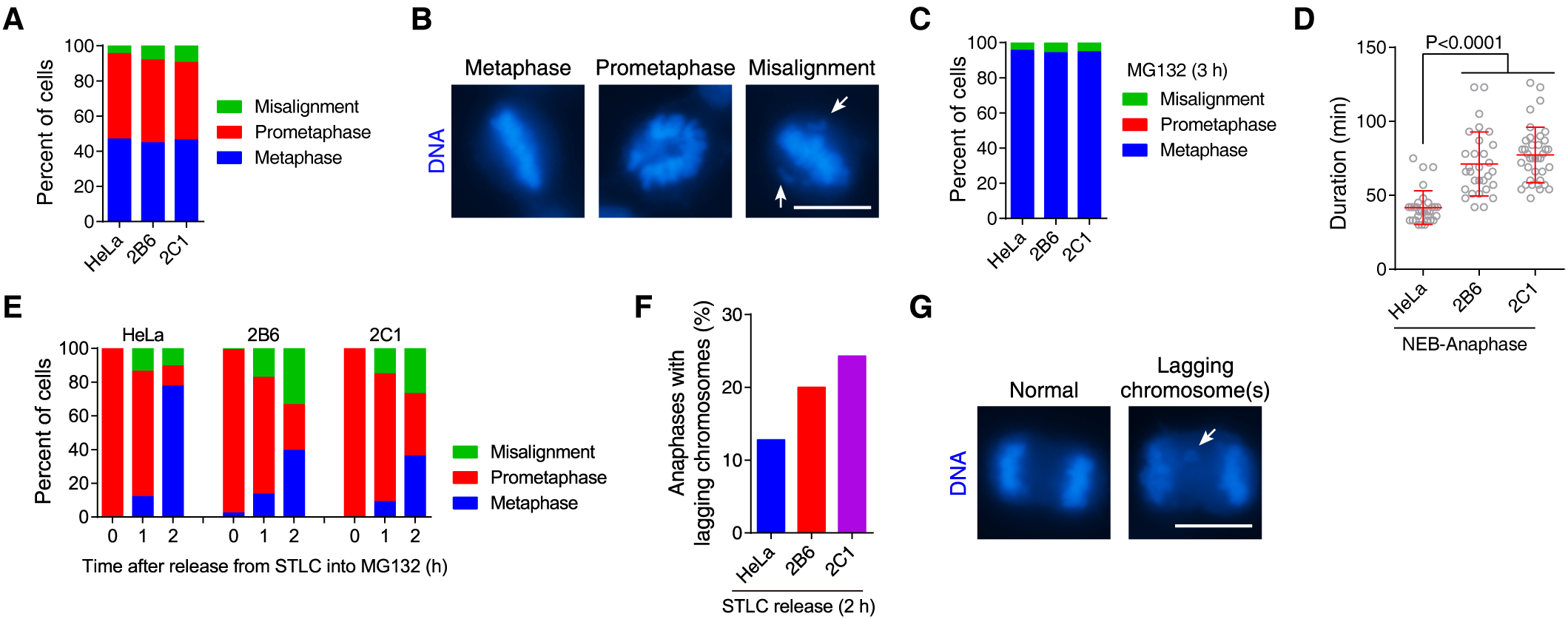
Loss of Bub1 does not compromise chromosome segregation fidelity during unperturbed mitosis. (A and B) Asynchronously growing HeLa cells and the indicated ΔBub1 clones were fixed for DNA staining. Preanaphase mitosis was classified and quantified in 100 cells for each condition (A). Example images are shown (B). (C) Asynchronously growing HeLa cells and the indicated ΔBub1 clones were exposed to MG132 for 3 h and then fixed for DAPI staining. Pre-anaphase mitosis was classified and quantified in 100 cells for each condition. (D) HeLa and the indicated ΔBub1 clones were released from double thymidine treatment, and then mitosis progression was analyzed by time-lapse live imaging of chromosomes labelled with a cell permeable fluorescent probe SiR-Hoechst (0.25 μM). The time from NEB to anaphase onset was determined in ≥30 cells. Means and standard deviations (SDs) are shown. (E) HeLa cells and the indicated ΔBub1 clones were exposed to STLC for 5 h, then washed into fresh medium with MG132 and fixed at the indicated time points. Approximately 100 cells were classified and quantified for each condition. (F and G) HeLa cells and the indicated ΔBub1 clones were released from 5 h treatment with STLC, then fixed at 2 h post-release and stained with DAPI. The percentage of cells with lagging chromosomes was determined in 100 anaphase cells for each condition (F). Example images are shown (G). Arrows point to misaligned chromosomes (B) and lagging chromosomes (G). Scale bars, 10 μm.

**Figure 3.**
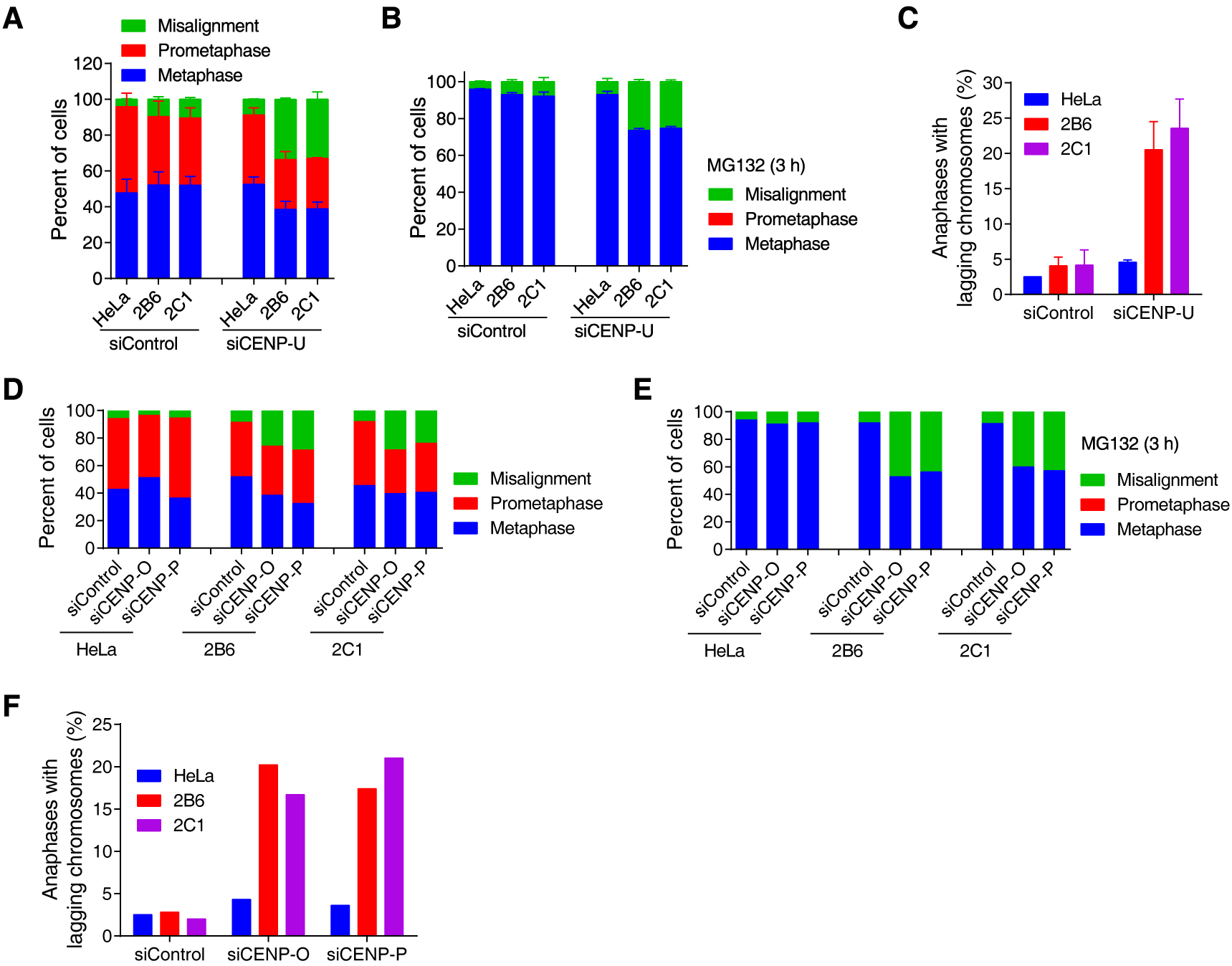
Knockdown of the CENP-U complex strongly affects chromosome alignment and segregation in ΔBub1 cells, but not in control HeLa cells. (A and B) HeLa cells and the indicated ΔBub1 clones were transfected with control siRNA and CENP-U siRNA, and then fixed for DNA staining (A), or were exposed to MG132 for 3 h prior to fixation (B). Pre-anaphase mitosis was classified and quantified in 300 cells from three independent experiments. Means and SDs are shown. (C) HeLa cells and the indicated ΔBub1 clones were transfected with control siRNA and CENP-U siRNA, then fixed for DNA staining. The percentage of cells with lagging chromosomes was determined in 200 anaphase cells from two independent experiments. Means and ranges are shown. (D and E) HeLa cells and the indicated ΔBub1 clones were transfected with control siRNA and CENP-O or CENP-P siRNA, and then fixed for DNA staining (D), or were exposed to MG132 for 3 h prior to fixation (E). Pre-anaphase mitosis was classified and quantified in around 100 cells or each condition. (F) HeLa cells and the indicated ΔBub1 clones were transfected with control siRNA and CENP-O or CENP-P siRNA, then fixed for DNA staining. The percentage of cells with lagging chromosomes was determined in 100 anaphase cells for each condition. See also Figure S2.

We next used time-lapse live imaging to examine mitosis progression in ΔBub1 cells. The time required for the transition from nuclear envelope breakdown (NEB) to anaphase onset was significantly increased in ΔBub1 cells (71.2 ± 4.0 min and 77.26 ± 3.1 min), compared with that in control HeLa cells (41.7 ± 2.1 min) (Figure 2D). The prolonged mitosis in ΔBub1 cells was due to the delay in the congression of one or a few chromosomes. We found that all chromosomes were eventually able to align on the metaphase plate, and that whole chromosome missegregation was rarely observed in ΔBub1 cells, indicating that the SAC functionally operated.

We further examined chromosome congression and segregation when cells were released from transient mitotic arrest induced by S-trityl-L-cysteine (STLC) (Lampson *et al*, 2004), which is an Eg5 inhibitor that prevents centrosome separation during mitotic entry and causes the formation of monopolar spindles with erroneously attached chromosomes (Skoufias *et al*, 2006). We treated cells with STLC to accumulate monopolar mitoses and then released them into MG132 to allow bipolar spindle formation and chromosome alignment. We found that, at 2 h post-release from STLC, while 77.6% control HeLa cells have completed metaphase chromosome alignment, only 36.2%-39.9% ΔBub1 cells have done this (Figure 2E). Consistently, when cells were released from STLC into fresh medium without MG132, the percentage of anaphase cells with lagging chromosomes increased from 12.8% in control HeLa cells to 20%-24.3% in ΔBub1 cells (Figure 2F and 2G). Thus, upon release from transient mitotic arrest during which erroneous attachments were generated, ΔBub1 cells were impaired in correcting erroneous KT-MT attachments.

Thus, loss of Bub1 does not cause obvious defects in whole chromosome segregation during unperturbed mitosis.

### Knockdown of the CENP-U complex strongly affects chromosome alignment and segregation in ΔBub1 cells, but not in control HeLa cells

We then asked why loss of Bub1 does not compromise the fidelity of chromosome segregation in ΔBub1 cells. One possibility is that another protein might compensate for the loss of Bub1 in chromosome congression. Such a protein could play a role in the centromere/kinetochore localization of Aurora B and/or Plk1, which has a well-established role in chromosome bi-orientation (Carmena *et al*, 2012, Hindriksen *et al*, 2017, Saurin, 2018).

Centromere localization of Aurora B depends on the recruitment of the CPC by Bub1-mediated H2ApT120 and Haspin-mediated histone H3 threonine 3 phosphorylation (H3pT3) (Kelly *et al*, 2010, Wang *et al*, 2010, Yamagishi *et al*, 2010). We and others recently showed that simultaneous inhibition of H2ApT120 and H3pT3 did not obviously affect chromosome alignment (Broad *et al*, 2020, Hadders *et al*, 2020, Liang *et al*, 2020). This suggests that the H3pT3-dependent pool of Aurora B at centromeres may not play a role in supporting proper chromosome alignment in ΔBub1 cells.

The delay in congression of single or a few chromosomes in ΔBub1 cells is reminiscent of that observed in mitotic HeLa cells treated with compounds that inhibit the PBD-dependent interaction between Plk1 and its docking proteins (Reindl *et al*, 2008, Watanabe *et al*, 2009). Given that kinetochore localization of Plk1 was largely reduced in ΔBub1 cells (Figure 1F), and that CENP-U recruits Plk1 to kinetochores through interacting with PBD (Kang *et al*, 2006), we speculated that the inner kinetochore CENP-U complex might account for the accurate chromosome segregation in ΔBub1 cells.

We therefore examined the effect of RNA interference (RNAi)-mediated depletion of the CENP-U complex on chromosome alignment and segregation. Analysis of asynchronously growing cells showed that CENP-U depletion only marginally increased chromosome alignment in control HeLa cells, as well as in Bub1 mutant clones 2D1 and 2D4 (Figures 3A and S2A). Strikingly, the percentage of mitotic ΔBub1 cells with misaligned chromosomes was over 3-fold higher upon CENP-U RNAi (33%-33.6%) than control RNAi (9.7%-10.4%). Moreover, after treatment with MG132 for 3 h, much more ΔBub1 cells were defective in metaphase chromosome alignment upon CENP-U RNAi than upon control RNAi (Figures 3B and S2B). Similar results were observed in control HeLa and ΔBub1 clone 2B6 cells when CENP-U was depleted by another siRNA (Figure S2C and S2D). Consistently, CENP-U depleted ΔBub1 cells exhibited dramatically higher rate of anaphases with lagging chromosomes (20.5%-23.6%) than control RNAi ΔBub1 cells (4.1%-4.2%) (Figure 3C).

Similarly, RNAi of CENP-O and CENP-P strongly affected chromosome alignment and segregation in ΔBub1 cells, but not in control HeLa cells (Figures 3D-3F, and S2E). This is in agreement with the interdependent inner kinetochore localization of the CENP-U complex subunits (Hori *et al*, 2008). The efficacy of the siRNAs was confirmed by knocking down of exogenously expressed CENP-U, CENP-O and CENP-P (Figure S2F-S2H).

Collectively, these results demonstrate that ΔBub1 cells become strongly defective in chromosome congression and segregation upon loss of the non-essential CENP-U complex.

### Knockdown of the CENP-U complex in ΔBub1 cells reduces the kinetochore/centromere localization of Plk1 but not the CPC

We then investigated the mechanism by which the CENP-U complex promotes chromosome alignment in ΔBub1 cells. Given the interaction of CENP-U with Plk1 (Kang et al, 2006), we first assessed the contribution of the CENP-U complex to Plk1 localization at kinetochores. We found that CENP-U RNAi did not seem to affect centrosomal localization of Plk1 (Figure 4A). Strikingly, CENP-U RNAi strongly reduced the kinetochore localization of Plk1 in both HeLa cells and ΔBub1 cells (Figure 4A and 4B), indicating that CENP-U contributes to kinetochore accumulation of Plk1 independent of Bub1. The residual Plk1 at mitotic kinetochores in CENP-U depleted ΔBub1 cells could be due to partial depletion of CENP-U, or to the presence of additional Plk1 receptors at centromeres/kinetochores. Moreover, Knl1 localized normally to kinetochores in CENP-U depleted ΔBub1 cells (Figure 4C and 4D), indicating that assembly of the outer kinetochore was not altered in these cells. Thus, CENP-U is specifically required for the kinetochore localization of Plk1.

**Figure 4.**
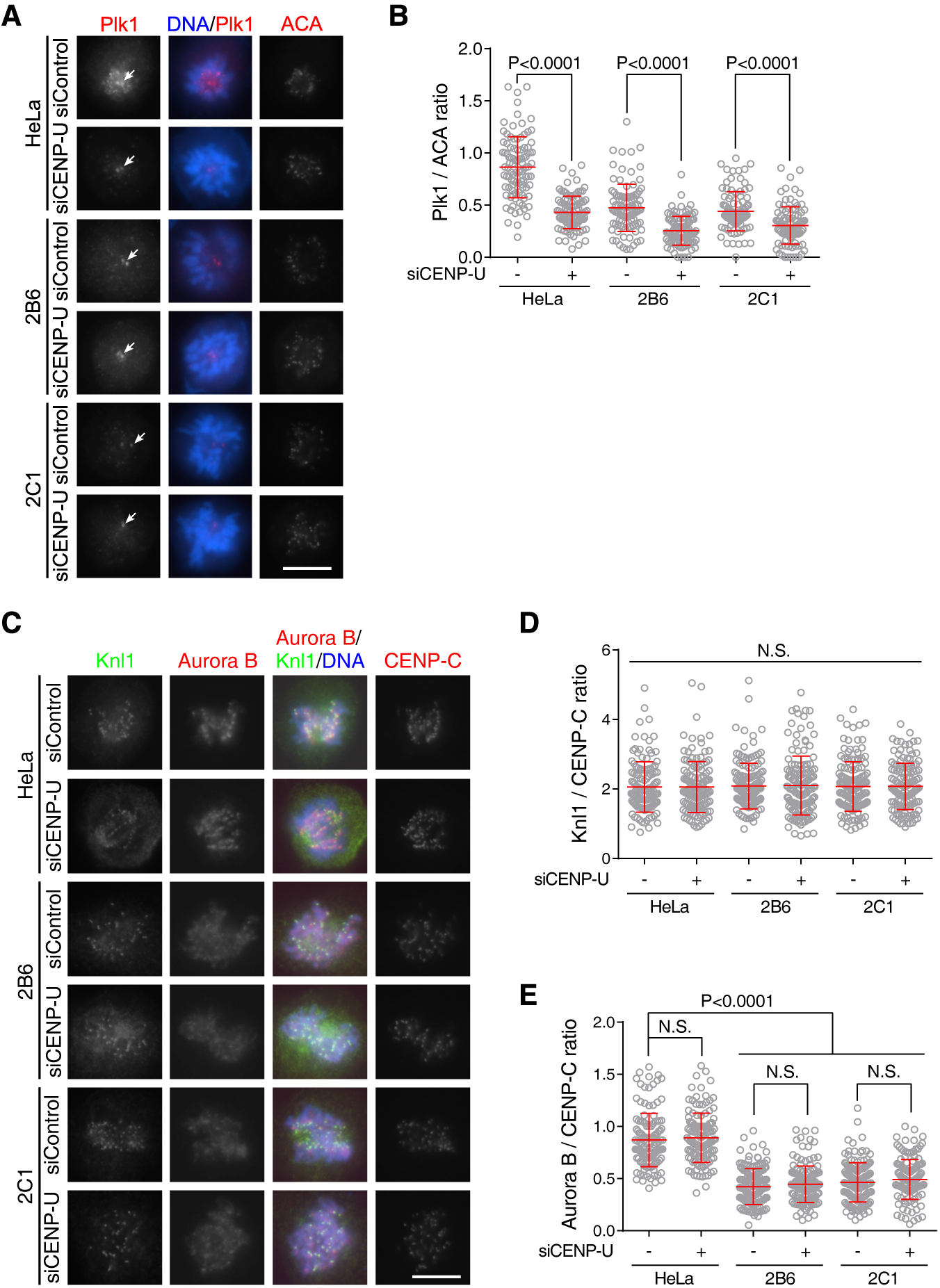
Knockdown of the CENP-U complex in ΔBub1 cells reduces the kinetochore/centromere localization of Plk1 but not the CPC. (A and B) HeLa cells and the indicated ΔBub1 clones were transfected with control siRNA and CENP-U siRNA, exposed to STLC for 5 h, and then immunostained with ACA and the Plk1 antibody. Example images are shown (A). The immunofluorescence intensity ratio of Plk1/ACA was determined in 20 cells (B). Arrows point to Plk1 that localized to centrosomes. (C-E) HeLa cells and the indicated ΔBub1 clones were transfected with control siRNA and CENP-U siRNA, and then immunostained with antibodies for Aurora B, CENP-C, and Knl1. Example images are shown (C). For each experiment, the immunofluorescence intensity ratios of Knl1/CENP-C (D) and Aurora B/CENP-C (E) were determined from 150 chromosomal regions containing >450 centromeres in 15 cells. Means and SDs (B, D and E) are shown. N.S., not significant. Scale bars, 10 μm. See also Figure S3.

It was recently reported that the budding yeast homolog of the CENP-U complex, the COMA complex composed of Ame1/Okp1^CENP-U/Q^ and Ctf19/Mcm21^CENP-P/O^, interacts with Sli15^INCENP^ *in vitro*, recruits Sli15/Ipl1^INCENP/Aurora B^ to the inner kinetochore, and promotes chromosome bi-orientation (Fischbock-Halwachs *et al*, 2019, Garcia-Rodriguez *et al*, 2019). We next examined whether the CENP-U complex contributes to the centromeric localization of the CPC. Quantitative immunofluorescence microscopy showed that, compared to that in control HeLa cells, the centromeric localization of Aurora B was reduced by 2-fold in ΔBub1 cells (Figure 4C and 4E). However, CENP-U RNAi did not detectably affect centromeric accumulation of Aurora B in either HeLa cells or ΔBub1 cells. Similarly, RNAi of CENP-U, CENP-P and CENP-O did not reduce the localization of Aurora B and INCENP at mitotic centromeres in HeLa cells (Figure S3). Thus, the CENP-U complex seems dispensable for centromeric localization of the CPC in HeLa cells.

Collectively, these results demonstrate that RNAi-mediated depletion of the CENP-U complex in ΔBub1 cells reduces the kinetochore/centromere localization of Plk1 but not the CPC.

### The CENP-U complex is sufficient to recruit Plk1, but not the CPC, independent of Bub1

To explore whether the CENP-U complex promotes chromosome congression in ΔBub1 cells through Plk1 and/or Aurora B, we next examined the capability of the CENP-U complex to recruit Plk1 and the CPC.

We first used an engineered U2OS cell line in which multiple copies of Lac Operator (LacO) repeats were stably integrated in an euchromatic region of chromosome 1 (Janicki *et al*, 2004). As a negative control, tethering EGFP-fused Lac repressor (EGFP-LacI) alone to the LacO array did not recruit Plk1 (Figure 5A). Remarkably, tethering CENP-U as a fusion protein with EGFP-LacI (EGFP-LacI-CENP-U) to the LacO array recruited Plk1 in around 60% of interphase cells. In contrast, tethering CENP-O, CENP-P, CENP-Q and CENP-R as EGFP-LacI fusion proteins did not recruit Plk1. These data imply that CENP-U directly interacts with Plk1. Indeed, the threonine to alanine mutation of residue 78 in CENP-U, whose phosphorylation is required for Plk1 binding (Kang *et al*, 2006), prevented EGFP-LacI-CENP-U from recruiting Plk1 (Figure 5B and 5C). Moreover, while EGFP-LacI-CENP-U recruited Plk1 to the LacO array in every mitotic cells we examined (Figure 5D), the EGFP-LacI-CENP-U-T78A mutant was unable to do so, indicating a critical requirement for T78 phosphorylation in binding Plk1.

**Figure 5.**
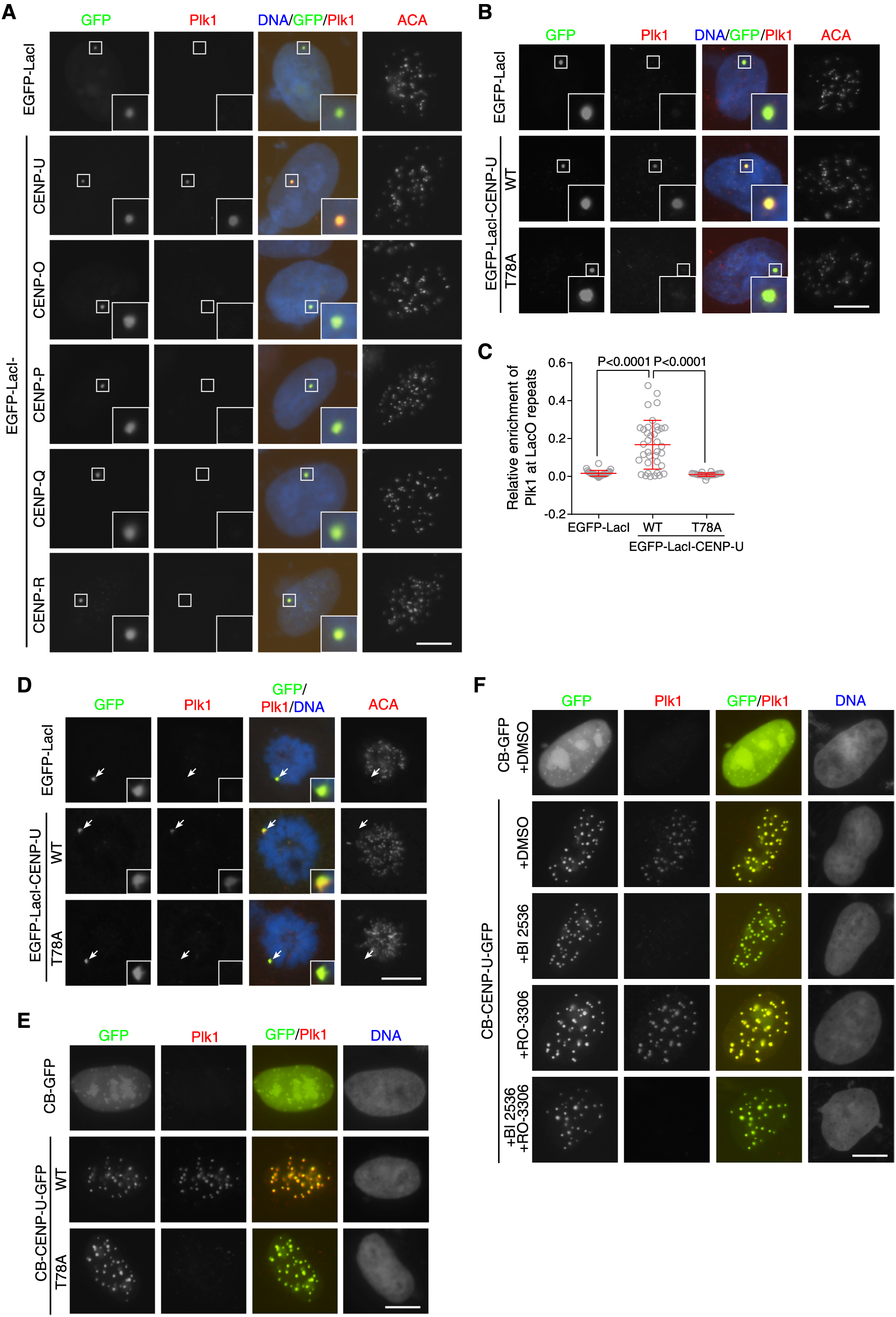
The CENP-U complex is sufficient to recruit Plk1, but not the CPC, independent of Bub1. (A) U2OS-LacO cells expressing EGFP-LacI and EGFP-LacI-fused CENP-U, O, P, Q, R were fixed and then subjected to immunofluorescence staining with ACA and the Plk1 antibody. (B and C) U2OS-LacO cells expressing EGFP-LacI and EGFP-LacI-CENP-U (WT or T78A) were fixed and then subjected to immunofluorescence staining with ACA and antibodies for GFP and Plk1. Example images of interphase cells are shown (B). The relative enrichment of Plk1 at the LacO repeats was quantified in 22-37 cells (C). Means and SDs are shown. (D) U2OS-LacO cells expressing EGFP-LacI and EGFP-LacI-CENP-U (WT or T78A) were exposed to 100 ng/ml nocodazole for 6 h. Mitotic cells were cytospun onto coverslips, fixed, and subjected to immunofluorescence staining with ACA, and antibodies for GFP and Plk1. Arrows point to the transgene loci. (E) HeLa cells expressing CB-GFP and CB-CENP-U-GFP (WT or T78A) were subjected to immunofluorescence staining with the Plk1 antibody. (F) HeLa cells expressing CB-GFP and CB-CENP-U-GFP were treated with 100 nM BI 2536, 9 μM RO-3306 or DMSO as control for 3 h, then subjected to immunofluorescence staining with the Plk1 antibody. Scale bars, 10 μm. See also Figure S4.

We further assessed the capability of CENP-U to recruit Plk1 to centromeres. We expressed CENP-U as a fusion protein with the centromere DNA-binding domain of CENP-B and GFP (CB-CENP-U-GFP) in HeLa cells, using CB-GFP as a negative control. Strikingly, centromere-tethered CENP-U, but not the CENP-U-T78A mutant, was capable of recruiting Plk1 (Figure 5E). Treatment with the Plk1 inhibitor BI 2536 prevented CB-CENP-U-GFP from recruiting Plk1 (Figure 5F). In contrast, treatment with the Cdk1 inhibitor RO-3306 did not affect the recruitment of Plk1 by CB-CENP-U-GFP, indicating that Cdk1 kinase activity is not critically required for the CENP-U-Plk1 interaction.

We next examined whether the CENP-U complex is able to recruit the CPC in human cells. As a positive control, tethering Haspin kinase as a fusion protein with EGFP-LacI (EGFP-LacI-Haspin) to the LacO array recruited the INCENP subunit of the CPC (Figure S4), presumably through the binding of CPC with Haspin-generated H3pT3 (Liang et al, 2020). In contrast, tethering CENP-O, CENP-P, CENP-Q, CENP-U and CENP-R as EGFP-LacI fusion proteins did not recruit INCENP. Thus, the CENP-U complex tethered to the LacO array is incapable of recruiting the CPC.

These results indicate that the CENP-U complex is sufficient to recruit Plk1, but not the CPC. Thus, it is unlikely that the CENP-U complex regulates chromosome alignment and segregation via binding the CPC as proposed in budding yeast (Fischbock-Halwachs *et al*, 2019, Garcia-Rodriguez *et al*, 2019). Since presumably Bub1 is not present either at the LacO transgene array or at interphase centromeres, we conclude that CENP-U is sufficient to recruit Plk1 in a manner independent of Bub1.

### Bub1 is sufficient and necessary for the recruitment of Plk1 to kinetochores independent of CENP-U

Having addressed the Bub1-independent role for CENP-U in recruiting Plk1, we next examined whether Bub1 can regulate Plk1 localization independent of CENP-U. As mentioned above, Plk1 was strongly reduced at kinetochores in ΔBub1 cells (Figure 1F). We further found that exogenous expression of GFP-fused Bub1 (Bub1-GFP) in ΔBub1 cells restored proper kinetochore localization of Plk1 (Figure 6A and 6B). However, this was not the case upon expression of the Bub1-T609A-GFP mutant, in which threonine 609 in the PBD-binding S-T_609_-P motif was mutated to alanine. These results indicate that Plk1 delocalization from kinetochores in ΔBub1 cells is due to specific loss of the Bub1-Plk1 interaction

**Figure 6.**
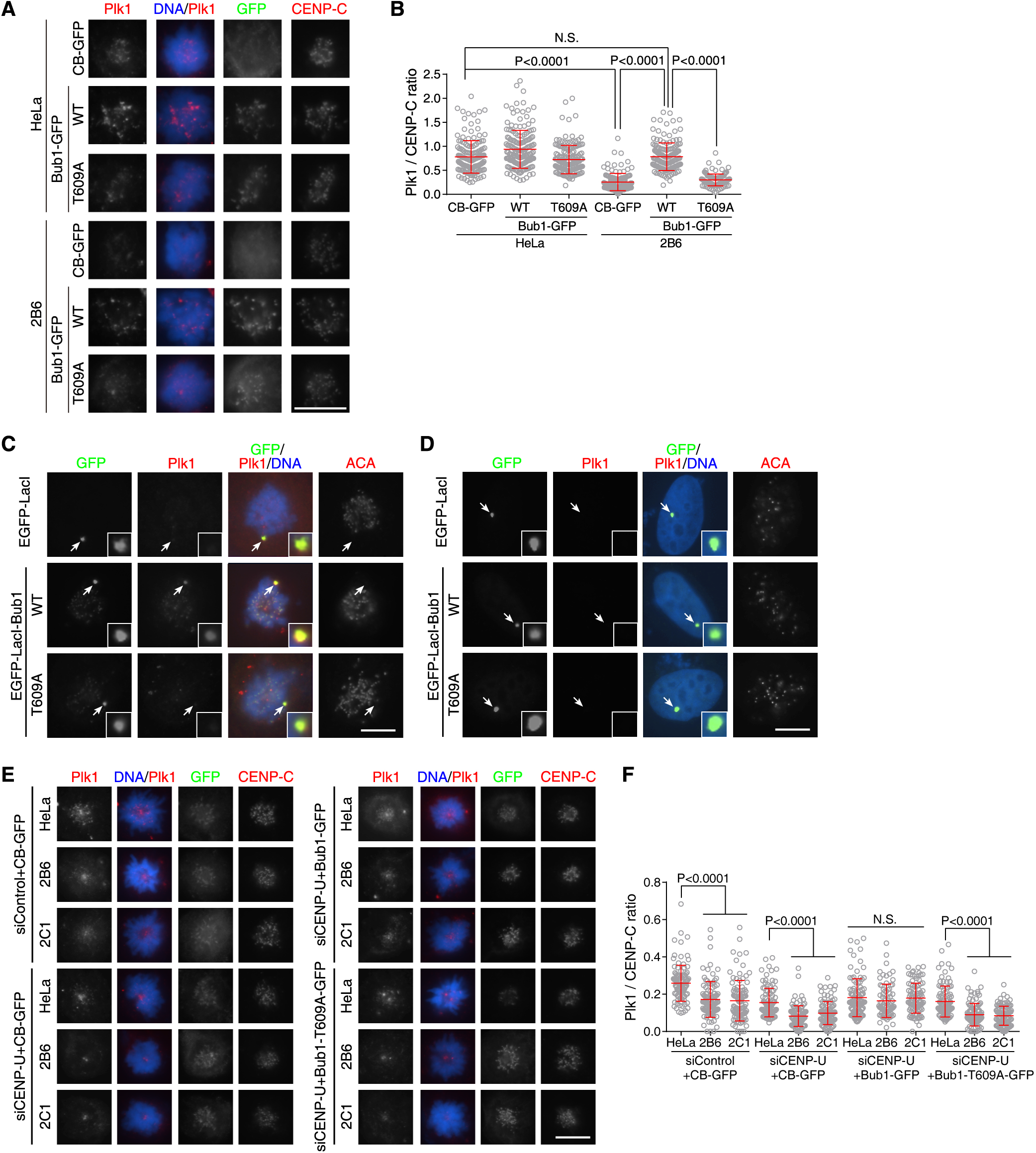
Bub1 is sufficient and necessary for the recruitment of Plk1 to kinetochores independent of CENP-U. (A) HeLa cells and ΔBub1 clone 2B6 were transfected with CB-GFP and Bub1-GFP (WT or 609A), exposed to 20 ng/ml nocodazole for 4 h, and then immunostained with antibodies for CENP-C, GFP and Plk1. Example images are shown (A). The immunofluorescence intensity ratio of Plk1/CENP-C was determined from 150 chromosomal regions containing >450 centromeres in 15 cells (B). (C) U2OS-LacO cells expressing EGFP-LacI and EGFP-LacI-Bub1 (WT or T609A) were exposed to 100ng/ml nocodazole for 6 h. Mitotic cells were cytospun onto coverslips, fixed, and subjected to immunofluorescence staining with ACA, and antibodies for GFP and Plk1. Arrows point to the transgene loci. (D) U2OS-LacO cells expressing EGFP-LacI and EGFP-LacI-Bub1 (WT or T609A) were fixed and then subjected to immunofluorescence staining with ACA and antibodies for GFP and Plk1. Arrows point to the transgene loci. (E and F) HeLa cells and the indicated ΔBub1 clones were transfected with control siRNA or CENP-U siRNA, together with expressing CB-GFP and Bub1-GFP (WT or T609A), exposed to STLC for 5 h, and then immunostained with antibodies for CENP-C, GFP and Plk1. Example images are shown (E). The immunofluorescence intensity ratio of Plk1/CENP-C was determined from 100 chromosomal regions in 20 cells (F). Means and SDs are shown (B and F). N.S., not significant. Scale bars, 10 μm.

We next examined whether Bub1 is sufficient to recruit Plk1, using the U2OS-LacO cell line. Immunofluorescence microscopy showed that tethering EGFP-LacI-fused Bub1 (EGFP-LacI-Bub1), but not the EGFP-LacI-Bub1-T609A mutant, recruited Plk1 to the LacO array in mitosis (Figure 6C). In contrast, EGFP-LacI-Bub1 could not recruit Plk1 to the LacO array in interphase (Figure 6D), likely due to loss of Cdk1 phosphorylation of the PBD-binding S-T_609_-P motif (Qi *et al*, 2006). Since presumably the CENP-U complex was absent at the LacO array, these results demonstrated that Bub1 is sufficient to recruit Plk1 during mitosis independent of CENP-U.

We then examined whether Bub1 can support the kinetochore localization of Plk1 in CENP-U depleted cells. For this purpose, we transfected HeLa and ΔBub1 cells with control siRNA or CENP-U siRNA, then expressed Bub1 - GFP and the Bub1-T609A-GFP mutant, using CB-GFP as the negative control. Immunofluorescence microscopy showed that, as expected, CENP-U knockdown strongly reduced the kinetochore localization of Plk1 in HeLa and ΔBub1 cells expressing CB-GFP. Remarkably, expression of Bub1-GFP, but not the Bub1-T609A-GFP mutant, restored the kinetochore localization of Plk1 in ΔBub1 cells to a level comparable to that in CENP-U-depleted HeLa cells (Figure 6E and 6F). Thus, replacement of endogenous Bub1 with exogenously expressed Bub1-GFP is able to support Plk1 localization at the outer kinetochore in CENP-U depleted cells.

These data indicate that Bub1 is sufficient and necessary for the recruitment of Plk1 to the outer kinetochore during mitosis, in a CENP-U-independent manner.

### Bub1-mediated kinetochore localization of Plk1 is essential for proper chromosome alignment in CENP-U depleted cells

Based on all of the data above, we speculated that loss of Bub1-mediated localization of Plk1 at the outer kinetochore underlies the chromosome congression defects in ΔBub1 cells depleted of CENP-U. If so, restoring proper accumulation of Plk1 at the outer kinetochore by exogenous expression of Bub1 should be able to rescue the chromosome alignment defect in these cells.

To test this, we transfected HeLa and ΔBub1 cells with control siRNA or CENP-U siRNA, then expressed Bub1 - GFP and the Bub1-T609A-GFP mutant, using CB-GFP as the negative control. Cells were arrested in mitosis by STLC treatment, and then released into MG132 for 3 h to allow bipolar spindle formation and chromosome alignment, as previously described (Lampson *et al*, 2004). As expected, CENP-U knockdown caused a strong defect in chromosome alignment in ΔBub1 cells expressing CB-GFP. Importantly, expression of Bub1-GFP, but not Bub1-T609A-GFP, largely restored proper chromosome alignment in CENP-U depleted ΔBub1 cells (Figure 7A-7C). Therefore, replacement of endogenous Bub1 with exogenously expressed Bub1-GFP in CENP-U depleted cells is able to support kinetochore localization of Plk1 and proper chromosome alignment.

**Figure 7.**
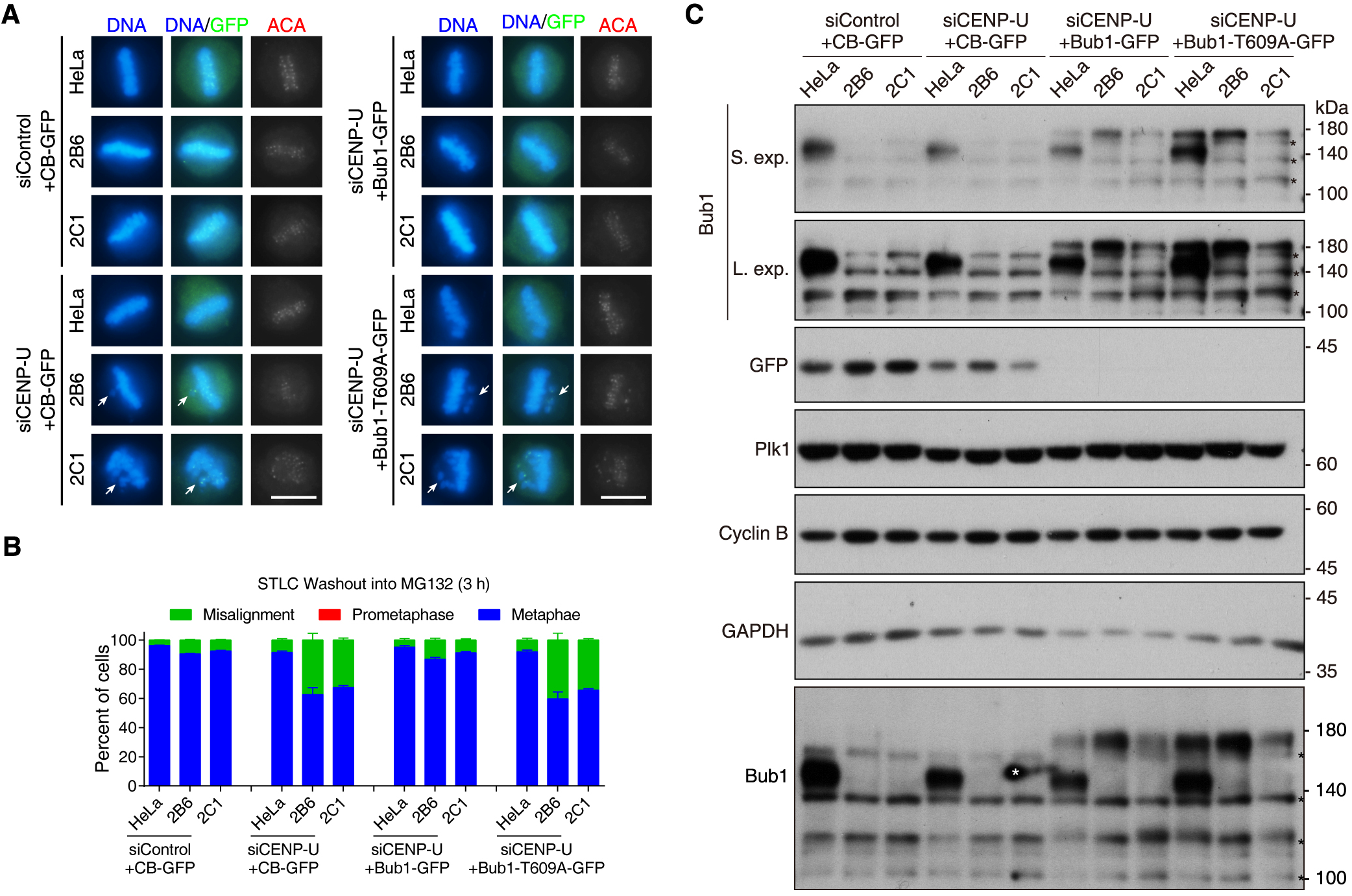
Bub1-mediated kinetochore localization of Plk1 is essential for proper chromosome alignment in CENP-U depleted cells. (A and B) HeLa cells and the indicated ΔBub1 clones were transfected with control siRNA or CENP-U siRNA, together with expressing CB-GFP and Bub1-GFP (WT or T609A), exposed to STLC for 5 h, then released into fresh medium with MG132 and fixed at 3 h later for DAPI, GFP and ACA staining. Example images are shown (A). Arrows point to the misaligned chromosomes. Approximately 200 cells were classified and quantified for each condition from two independent experiments (B). Means and ranges are shown. (C) HeLa cells and the indicated ΔBub1 clones were transfected as in (A). Cells were treated with 300 ng/ml nocodazole for 15 h, and then mitotic cells were collected. The samples were analyzed by immunoblotting the indicated antibodies and rabbit anti-Bub1 polyclonal antibodies. * represents non-specific bands or contaminated background signal. N.S., not significant. S.exp., short exposure; L.exp., long exposure. Scale bars, 10 μm.

We therefore conclude that Bub1-mediated kinetochore localization of Plk1 is essential for proper chromosome alignment in the absence of CENP-U.

## DISCUSSION

Bub1 is a highly conserved multi-task protein kinase, which has long been considered important for chromosome congression and SAC signaling during mitosis (Bolanos-Garcia & Blundell, 2011, Johnson *et al*, 2004, Perera *et al*, 2007, Taylor & McKeon, 1997). However, recent studies report surprising evidence that Bub1 may not be essential in human cells (Currie *et al*, 2018, Raaijmakers & Medema, 2019, Raaijmakers *et al*, 2018, Rodriguez-Rodriguez *et al*, 2018, Zhang *et al*, 2019a, Zhang *et al*, 2019b), with the underlying mechanism unknown.

To analyze the role for Bub1 kinase in mitosis, we attempted to knock out Bub1 in HeLa cells using CRISPR/Cas9 with a small guide (sg)RNA targeting genome DNA in exon 20 which encodes residues 793-800 in the kinase domain (residues 784-1085). We obtained a number of clonal cell lines in which H2ApT120 is undetectable whereas the Bub1 protein level varies among these clones. In this study, we mainly used ΔBub1 clones 2B6 and 2C1 for functional analysis. Lack of Bub1 protein in these two clones is confirmed by immunoblotting and immunofluorescence microscopy using two polyclonal antibodies, generated by immunization of recombinant GST-Bub1 (residues 336-489) in sheep and synthesized peptide (residues 536-551) in rabbit (Taylor *et al*, 2001, Zhang *et al*, 2019c). We cannot fully rule out the possibility for the residual expression of kinase-deficient truncated and/or internally deleted mutant forms of Bub1 that lack the epitopes recognized by the antibodies we used (Rodriguez-Rodriguez *et al*, 2018). Importantly, compared to Bub1 mutant clones 2D1 and 2D4 which expressing residual Bub1, ΔBub1 clones 2B6 and 2C1 show drastically high sensitivity upon knockdown of the CNEP-U complex.

In this study, using ΔBub1 clones 2B6 and 2C1, we show that Bub1 plays a redundant role with the non-essential CENP-U complex in recruiting Plk1 to the kinetochore. While loss of either pathway of Plk1 recruitment does not compromise the accuracy of whole chromosome segregation, disrupting both pathways causes a remarkable decrease in the kinetochore localization of Plk1 under a threshold level that is required for proper chromosome alignment and segregation. Thus, parallel recruitment of Plk1 to kinetochores by Bub1 and the CENP-U complex ensures the fidelity of mitotic chromosome segregation. This study provides important insight into the molecular mechanism for the non-essentiality of Bub1, which may have implications for targeted treatment of cancer cells harboring mutations in either Bub1 or the CENP-U complex.

A number of proteins, including CENP-U and Bub1, are reported to be Plk1 receptors at centromere/kinetochore (Combes *et al*, 2017, Lens *et al*, 2010, Lera *et al*, 2016, Singh *et al*, 2021). The functional significance for the role of CENP-U and Bub1 in recruiting Plk1 to kinetochores has been elusive (Kang *et al*, 2006, Qi *et al*, 2006). Our findings indicate that these two recruitment pathways are redundant in ensuring that Plk1 is kinetochore-localized for proper chromosome alignment and segregation in mitosis. Our conclusion is supported by a manuscript recently deposited by Iain Cheeseman Lab in bioRxiv (DOI: 10.1101/2020.11.30.404327), in which the authors investigated the source for the differential requirement of the CENP-U complex across human cell lines and revealed parallel Plk1 kinetochore recruitment pathways.

While this manuscript was under preparation, a study reported how Cdk1 and Plk1 promote kinetochore recruitment of Plk1 onto Bub1 and CENP-U (Singh *et al*, 2021). Singh et al. mainly used in vitro reconstituted kinetochore to report that binding of Plk1 to CENP-U is strictly dependent on priming Cdk1 activity, which phosphorylates threonine 98 to prime Plk1 binding and, in turn, enhances threonine 78 phosphorylation to create a stable docking configuration. Our observations shown in Figure 5 indicate that, at least when tethered to the LacO transgene array or to centromeres, Cdk1 kinase activity does not play a crucial role in CENP-U-mediated recruitment of Plk1 in cells. Our work extends these studies, and most importantly, reveals the functional significance of Plk1 recruitment by CENP-U at the inner kinetochore.

Plk1 localizes to various locations during mitosis, including centrosomes, spindle, centromere/kinetochore, central spindle, and midbody (Petronczki *et al*, 2008). Plk1 plays multiple roles in mitotic entry, bi-polar spindle formation, chromosome alignment, SAC signaling, cohesin removal from chromosome arms, and cytokinesis (Combes *et al*, 2017). RNAi and kinase inhibitors are commonly used to dissect the functions of Plk1 (Lenart *et al*, 2007, Steegmaier *et al*, 2007, Strebhardt, 2010). These approaches have the limitation to selectively remove Plk1 from specific locations. Given that we can now selectively remove Plk1 from kinetochores, future studies are required to dissect the molecular mechanisms by which kinetochore-localized Plk1 promotes chromosome alignment and centromere protection (Addis Jones *et al*, 2019, Lens *et al*, 2010, Lera *et al*, 2019, Olukoga *et al*, 2019).

## ACKNOWLEDGEMENTS

We thank Drs. David Spector and Stephen Taylor for kindly providing reagents; and the Life Sciences Institute core facility for technical assistance. This work was supported by grants to F. Wang from the National Science Fund for Distinguished Young Scholars (32025011), National Natural Science Foundation of China (31771499, 32061160470, 31571393, 31322032, 31371359, and 31561130155), the National Key Research and Development Program of China (2017YFA0503600), the Natural Science Foundation of Zhejiang Province (LZ19C070001), and the Royal Society Newton Advanced Fellowship (NA140075).

## AUTHOR CONTRIBUTIONS

Q.C., M.Z., and X.P. designed and performed the majority of the experiments and analyzed the data, with contributions from L.Z, and H.Y. F.W. conceived the project, designed the experiments, analyzed the data, and wrote the manuscript.

## SUPPLEMENTAL FIGURES AND LEGENDS

**Figure S1.**
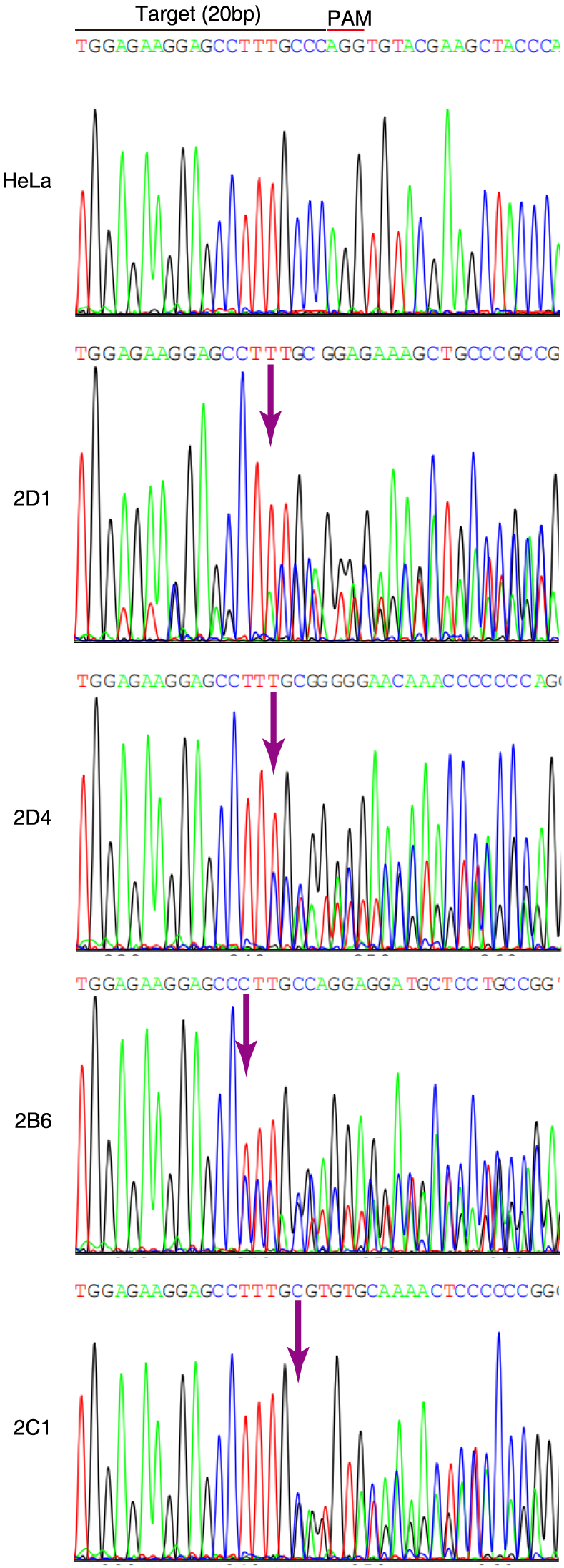
Knockout of Bub1 by CRISPR/Cas9 in HeLa cells (Related to Figure 1) Genomic DNA sequencing of control HeLa cells and the Bub1 mutant clones. The genomic DNA PCR fragments were sequenced directly. The sgRNA target DNA sequence preceding a 5-NGG protospacer adjacent motif (PAM) is shown. Arrows point to indels.

**Figure S2.**
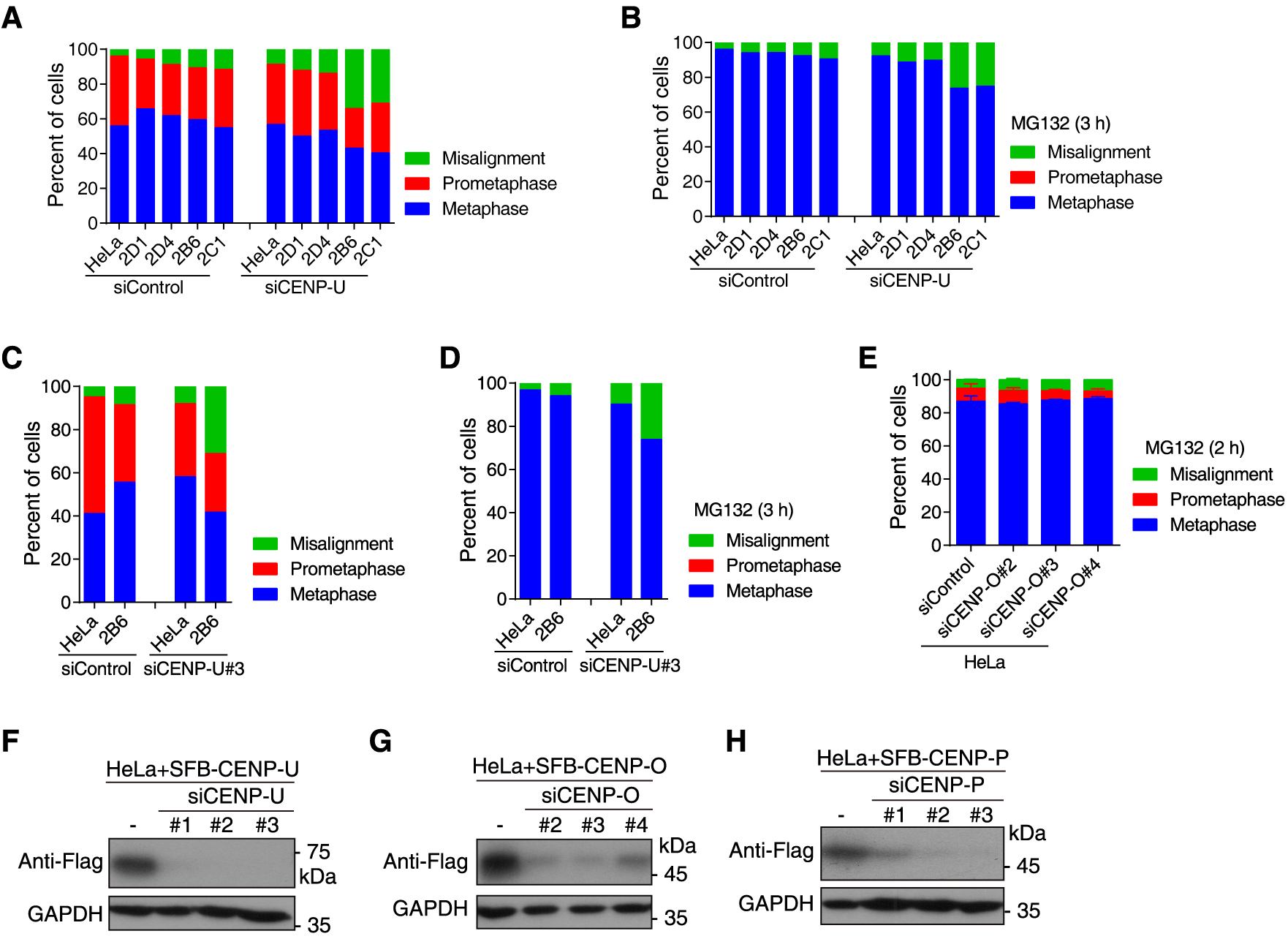
Knockdown of the CENP-U complex strongly affects chromosome alignment and segregation in ΔBub1 cells, but not in control HeLa cells (Related to Figure 3) (A) HeLa cells and the indicated Bub1 mutant clones were transfected with control siRNA and CENP-U siRNA, and then fixed for DNA staining. Pre-anaphase mitosis was classified and quantified in 100 cells. (B) HeLa cells and the indicated Bub1 mutant clones were transfected with control siRNA and CENP-U siRNA, and then exposed to MG132 for 3 h prior to fixation. Pre-anaphase mitosis was classified and quantified in 100 cells. (C) HeLa cells and the indicated ΔBub1 clones were transfected with control siRNA and CENP-U siRNA#3, and then fixed for DNA staining. Pre-anaphase mitosis was classified and quantified in 100 cells. (D) HeLa cells and the indicated ΔBub1 clones were transfected with control siRNA and CENP-U siRNA#3, and then exposed to MG132 for 3 h prior to fixation. Pre-anaphase mitosis was classified and quantified in 100 cells. (E) HeLa cells were transfected with control siRNA and CENP-O siRNAs, and then exposed to MG132 for 2 h prior to fixation. Pre-anaphase mitosis was classified and quantified in 200 cells for each condition from two independent experiments. Means and ranges are shown. (F) HeLa cells expressing SFB tag-fused CENP-U were transfected with control siRNA and CENP-U siRNAs, and then lysed for immunoblotting with antibodies for the Flag tag and GAPDH. (G) HeLa cells expressing SFB-CENP-O were transfected with control siRNA and CENP-O siRNAs, and then lysed for immunoblotting with antibodies for the Flag tag and GAPDH. (F) HeLa cells expressing SFB-CENP-P were transfected with control siRNA and CENP-P siRNAs, and then lysed for immunoblotting with antibodies for the Flag tag and GAPDH.

**Figure S3.**
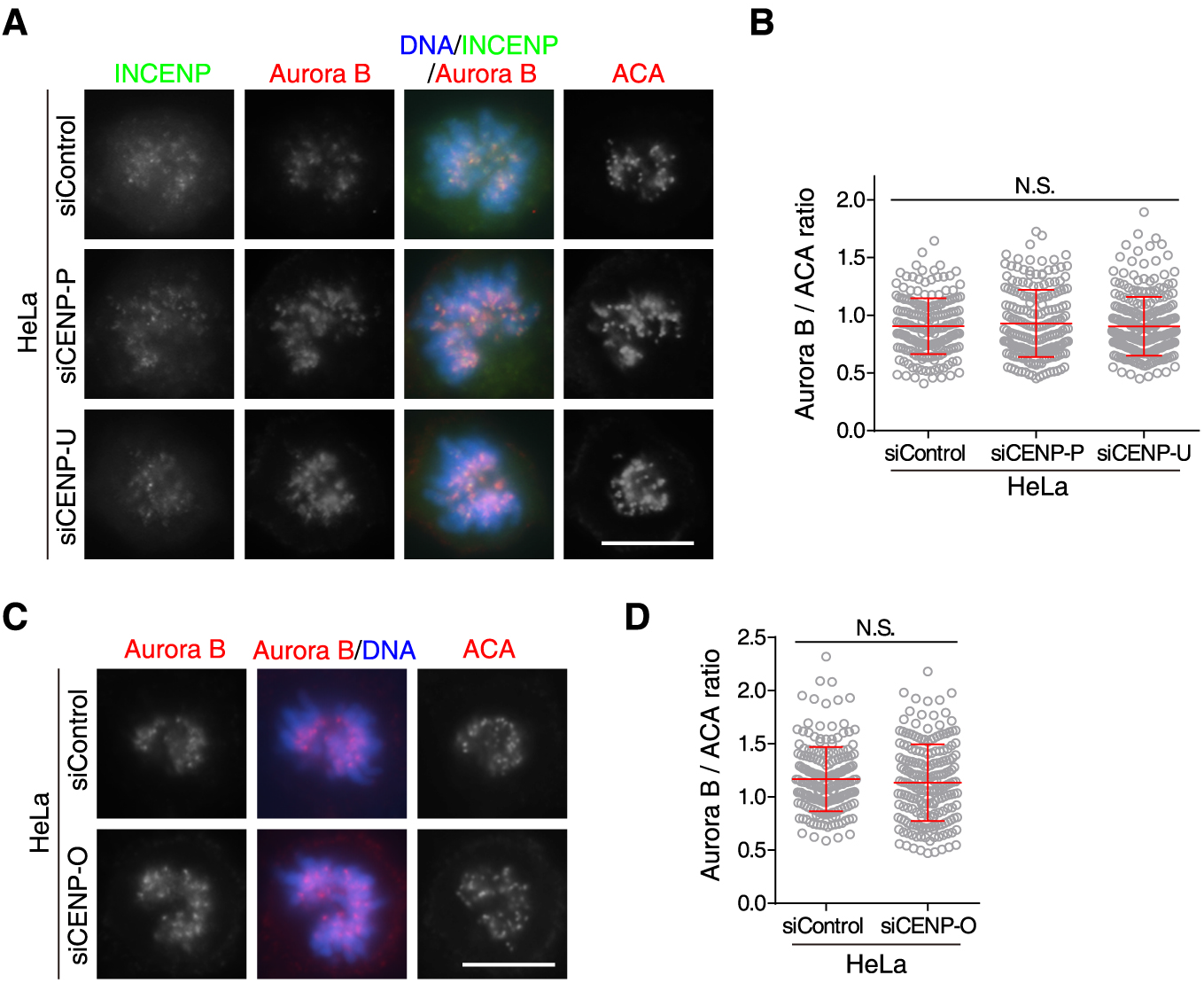
Knockdown of the CENP-U complex in ΔBub1 cells reduces the kinetochore/centromere localization of Plk1 but not the CPC (Related to Figure 4) (A and B) HeLa cells were transfected with control siRNA, CENP-P siRNA, or CENP-U siRNA, and then immunostained with ACA and antibodies for Aurora B and INCENP. Example images are shown (A). The immunofluorescence intensity ratio of Aurora B/ACA was determined from 200 chromosomal regions containing >600 centromeres in 20 cells (B). (C and D) HeLa cells were transfected with control siRNA or CENP-O siRNA, and then immunostained with ACA and the Aurora B antibody. Example images are shown (C). The immunofluorescence intensity ratio of Aurora B/ACA was determined from 200 chromosomal regions containing >600 centromeres in 20 cells (D). Means and SDs are shown (B and D). N.S., not significant. Scale bars, 10 μm.

**Figure S4.**
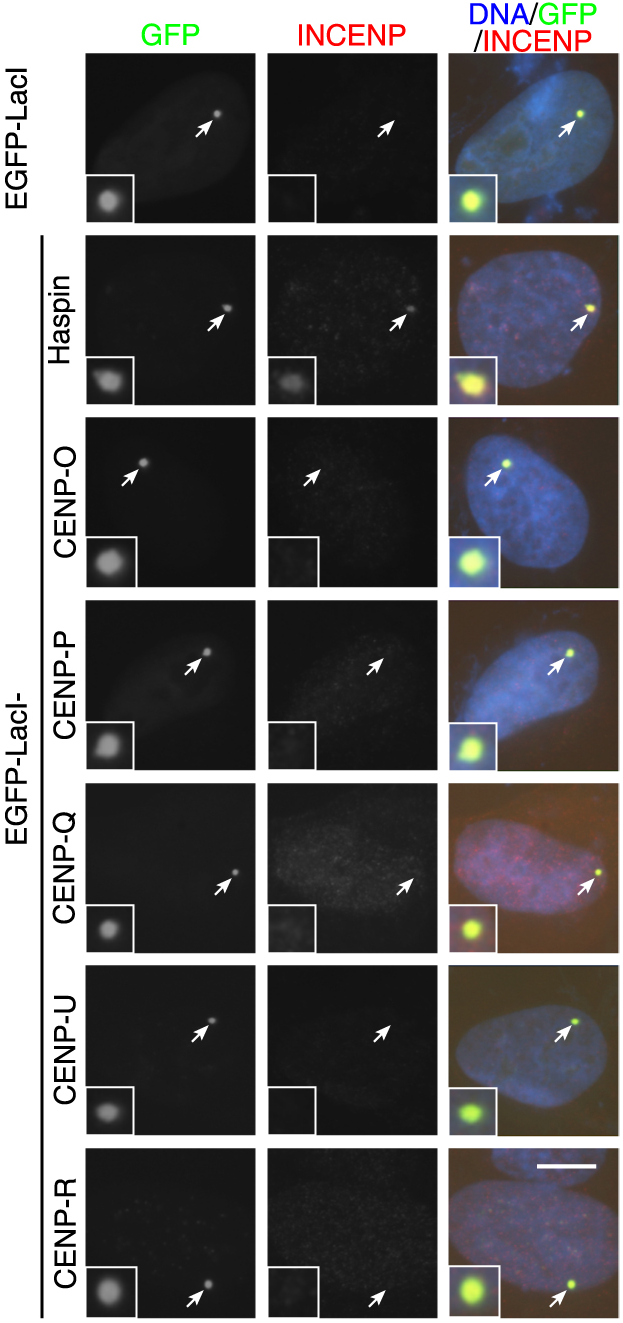
The CENP-U complex is not sufficient to recruit the CPC (Related to Figure 5) U2OS-LacO cells expressing EGFP-LacI, EGFP-LacI-Haspin, and EGFP-LacI-fused CENP-U, O, P, Q, R were fixed and then subjected to immunofluorescence staining with the INCENP antibody. Arrows point to the transgene loci. Scale bars, 10 μm.

## EXPERIMENTAL PROCEDURES

### Cell culture, plasmids, siRNA, transfection, and drug treatments

All cells were cultured in DMEM supplemented with 1% penicillin/streptomycin and 10% FBS (Gibco), and maintained at 37°C with 5% CO2. U2OS-LacO cells (kindly provided by Dr. David Spector) were maintained in the presence of 100 μg/ml hygromycin (Sigma).

To make the EGFP-LacI fusion constructs, the PCR fragments of Bub1, Haspin, and CENP-O, P, Q, U, R, were inserted into the BsiWI/BamHI sites of pSV2-EGFP-LacI. pBos-CENP-B-GFP was constructed by replacing the H2B fragment in pBos-H2B-GFP (Clontech) with the KpnI/BamHI-digested PCR fragments encoding the centromere-targeting domain (residues 1-163) of CENP-B. The pBos-Bub1-GFP construct was constructed by replacing the H2B fragment in pBos-H2B-GFP with the KpnI/BamHI digested PCR fragments encoding full-length Bub1. To make CENP-B-CENP-U-GFP constructs, the PCR fragments of CENP-U was inserted into the BamHI site of pBos-CENP-B-GFP. To make SFB (SFB is a triple tag of S tag, Flag tag, and streptavidin-binding peptide)-fused CENP-O, P, U constructs, the PCR-amplified DNA fragments of CENP-O, P, U were subcloned into the pDONR201 vector using Gateway Technology (Invitrogen). All point mutations were introduced with the QuikChange II XL site-directed mutagenesis kit (Agilent Technologies). All plasmids were sequenced to verify desired mutations and absence of unintended mutations.

The following siRNA duplexes ordered from Integrated DNA Technologies (IDT) or RiboBio were used: siCENP-U#1 (5’-GAAAGCCAUCUGCGAAAUAdTdT-3’, for Figure 3A-3C; Figure 4; Figure 6E, 6F; Figure 7; Figure S2A, S2B, S2F; Figure S3A, S3B), siCENP-U#2 (5’-GAAAAUAAGUACACAACGUdTdT-3’, for Figure S2F), siCENP-U#3 (5’-GGGAAGAUAUCUCAUGACAdTdT-3’, for Figure S2C, S2D, S2F), siCENP-O#2 (5’-GGACCUUGUCAUACAGAAAdTdT-3’, for Figure S2E, S2G), siCENP-O#3 (5’-GGAGCAACGAGCAUCUCAUdTdT-3’, for Figure 3D-3F; Figure S2E, S2G; Figure S3C, S3D), siCENP-O#4 (5’-GGACCUCACAGCAACUCUUdTdT-3’, for Figure S2E, S2G), siCENP-P#1 (5’-GAACCCUGGUAGGACUGCUUGGAAU-3’, for Figure 3D-3F; Figure S2H; Figure S3A, S3B), siCENP-P#2 (5’-UAUCGUAAGCGCACGUUUAdTdT-3’, for Figure S2H), siCENP-P#3 (5’-AGUCAUUGUUUGGAGGAUAdTdT-3’, for Figure S2H).

Plasmid and siRNA transfections were done with Fugene 6 (Promega) and Lipofectamine RNAiMAX (Invitrogen), respectively. Cells were arrested in S phase or at the G1/S boundary by single or double thymidine (2 mM, Calbiochem) treatment, respectively. Cells were arrested in prometaphase-like mitosis with 20-300 ng/ml nocodazole (Selleckchem), or STLC (5 μM, Tocris Bioscience), or were arrested in metaphase with MG132 (10 μM, Sigma). Mitotic cells were collected by selective detachment with “shake-off”.

### CRISPR/Cas9-mediated editing of Bub1 gene in HeLa cells

Single-guide RNA (sgRNA) for human Bub1 gene was ordered as oligonucleotides, annealed and cloned into the dual Cas9 and sgRNA expression vector pX330 (Addgene, #42230) with BbsI sites. The plasmids were transfected into HeLa cells using Fugene 6 (Promega) according to the manufacturer’s protocol. After 48 h incubation, the cells were split individually to make a clonal cell line with brief selection using 1 μg/ml puromycin for 2-3 days. To obtain mutant clones, cells with undetectable H2ApT120 as confirmed by immunostaining were isolated. The genomic DNA fragments were PCR amplified and sequenced to confirm the gene disruption.

### Antibodies and immunoblotting

Rabbit polyclonal antibodies used were to Mad1 (GTX105079-S, GeneTex), H2ApT120 (Active Motif), GFP (A11122, Invitrogen), Cyclin B1 (clone D5C10, CST), GAPDH (14C10, Cell Signaling Technology/CST), INCENP (P240, CST). Rabbit anti-Bub1 and anti-Knl1 polyclonal antibodies were produced by immunization with the synthetic peptides NYGLPQPKNKPTGAR and MDGVSSEANEENDNIERPVRRR, respectively. Mouse monoclonal antibodies used were to Aurora B (AIM-1; BD Biosciences), Bub3 (Clone 31, BD Transduction Laboratories), BubR1 (Dr. Jakob Nilsson), α-Tubulin (T-6047, Sigma), Sgo1 (3C11, Abnova), Plk1 (ab17057, Abcam), Flag-tag (M2, Sigma), TOP2A (M042-3, MBL). Sheep anti-Bub1 antibody was kindly provided by Dr. Stephen Taylor (University of Manchester, UK). Guinea pig polyclonal antibodies against CENP-C were from MBL (PD030). The anti-human centromere autoantibody (ACA) were from Immunovision. Secondary antibodies for immunoblotting were goat anti-rabbit or horse anti-mouse IgG-HRP (CST). Secondary antibodies for immunostaining were donkey anti-rabbit IgG-Alexa Fluor 488 or Cy3 (Jackson ImmunoResearch); anti-mouse IgG-Alexa Fluor 488 or 546 (Invitrogen) or Cy5 (Jackson ImmunoResearch); anti-human IgG-Alexa Fluor 647 (Jackson ImmunoResearch); Goat anti-guinea pig IgG-Alexa Fluor 647 (Invitrogen), anti-sheep IgG-Cy3 (Jackson ImmunoResearch).

SDS-PAGE and immunoblotting were carried out using standard procedures using whole cell lysates prepared in standard SDS sample buffer, or in P500 buffer containing 25 mM Tris-HCl, pH 7.5, 500 mM NaCl, 0.1% NP-40, 2 mM MgCl_2_, 10% glycerol, 1 mM dithiothreitol (DTT), protease inhibitor cocktail (P8340, Sigma), 1 mM PMSF, 0.1 μM okadaic acid (Calbiochem), 10 mM NaF and 20 mM β-glycerophosphate and benzonase (Merck).

### Fluorescence microscopy

Asynchronous U2OS-LacO cells were fixed with 2% PFA in PBS for 10 min and then extracted with 0.5% Triton X-100 for 5 min. For HeLa or ΔBub1 cells grown on coverslips, to stain sheep polyclonal anti-Bub1 antibodies (Figure 1B), Knl1(Figure 1E; Figure 4C), BubR1(Figure 1H), Mad1(Figure 1H), H2ApT120(Figure 1B, 1I, 1B), TOP2A(Figure 1I), INCENP(Figure S3A; Figure S4) and Aurora B(Figure 1J; Figure 4C; Figure S3A, S3C), cells were fixed with 2% PFA/PBS for 10 min, and then extracted by 0.5% Triton/PBS for 5 min. To stain Bub3(Figure 1G), cells were fixed with 2% PFA/PBS for 10 min, and then extracted by 0.5% Triton/PBS for 5 min, then fixed with 4% PFA in PBS for 10 min, then extracted with −20°C methanol for 5 min. To stain Plk1 (Figure 5A, 5B, 5D; Figure 6A, 6C, 6D, 6E), cells were fixed with 1% PFA/PBS 5min, and then treated with 0.1M glycine/PBS for 1 h, at last extracted by 0.1% Triton X-100/PBS for 3 min; Or fixed in 0.5% Triton-X-100 + 4% PFA in PBS for 10 minutes (Figure 1F; Figure 4A; Figure 5E, 5F). To stain rabbit anti-Bub1 polyclonal antibodies (Figure 1C), Cells were pre-extracted with 0.5% Triton in PHEM (60 mM Pipes, 25 mM Hepes, 10 mM EGTA, and 2 mM MgCl2, pH 6.9) for 5 min, then fixed with 4% PFA in PHEM for 20 min. For Figure 5D and Figure 6C, U2OS-LacO cells were treated with nocodazole (100 ng/ml) for 6 h. After attachment to glass coverslips by Cytospin at 1500 rpm for 5 min, cells were fixed with 1% PFA/PBS for 5 min, treated with 0.1M Glyine/PBS for 60 min, extracted with 0.5% Triton X-100/PBS for 3 min. Fixed cells were stained with primary antibodies for 1-2 h and secondary antibodies for 1 h, all with 3% BSA in PBS with 0.1%-0.5% Triton X-100 and at room temperature. DNA was stained for 10 min with DAPI. Fluorescence microscopy was carried out at room temperature using a Nikon ECLIPSE Ni microscope with a Plan Apo Fluor 60X Oil (NA 1.4) objective lens and a Clara CCD (Andor Technology).

Quantification of fluorescent intensity was carried out with ImageJ (NIH) using images obtained at identical illumination settings. Briefly, to quantify the relative intensity at the centromere/kinetochore, the average pixel intensity of antibody staining at centromeres/kinetochores, defined as a circular region encompassing at least three centromere pairs, was determined using ImageJ. After background correction, the ratio of centromere/kinetochore immunostaining intensity of these proteins versus that of ACA or CENP-C, was calculated for each circular region. To quantify the means and SDs for the relative intensity of proteins of interest at centromeres/kinetochores from individual experiments, a total of 200 circular regions from 20 cells (10 non-overlapping circular regions per cell) were plotted.

To quantify the relative enrichment of proteins of interest at the LacO transgene array in U2OS-LacO cells, the average pixel intensity of antibody staining, within circles encompassing the EGFP-LacI fusion protein fluorescent signal at LacO transgene array, and in the nearby nucleus, was determined. After background correction, the ratio of average immunostaining intensity at LacO repeats versus that in the nuclei was calculated.

### Time-lapse live cell imaging

Time-lapse live cell imaging was carried out with the GE DV Elite Applied Precision DeltaVision system (GE Healthcare) equipped with Olympus oil objectives of 60X (NA 1.42) Plan Apo N or 40X (NA 1.35) UApo/340 and an API Custom Scientific complementary metal-oxide semiconductor camera, and Resolve3D softWoRx imaging software. Cells were plated in four-chamber glass-bottomed 35-mm dishes (Cellvis) coated with poly-D-lysine, and filmed in a climate-controlled and humidified environment (37°C and 5% CO2). Chromosomes were labelled with a cell permeable fluorescent probe SiR-Hoechst (0.25 μM). Images were captured every 3 min. The acquired images were processed using Adobe Photoshop and Adobe Illustrator.

### Statistical analysis

Statistical analyses were performed with a two-tailed unpaired Student’s t test in GraphPad Prism 6. A p value of less than 0.05 was considered significant.

